# Stratifying cellular injury in Alzheimer’s disease by chaperonin containing TCP1 subunits 2 and 3

**DOI:** 10.64898/2026.05.10.724132

**Authors:** Jan Mulder, Tibor Hortobágyi, Tibor Harkany

## Abstract

Chaperonins complex into double-ringed octamers to aid peptide folding. Recent evidence implicates dysfunctional chaperonin subunits in cancer and neurodegenerative diseases because their deregulation exacerbates cellular injury. Nevertheless, a gap of knowledge exists regarding the expression and localization of chaperonin subunits in relation to amyloidogenic processes in Alzheimer’s disease (AD). Here, we show that reduced levels of chaperonin-containing TCP-1 subunits 2 (CCT2) and 3 (CCT3) stratify AD, with the subcellular distribution of their residua being mutually exclusive with both β-amyloid and hyperphosphorylated tau in neurons. We find CCT3 localized to a subset of glial fibrillary acidic protein-positive astrocytes in AD. Increased oxidative stress *in vitro* upregulated CCT3 expression in astrocyte-like U251 cells. Conversely, CCT3, but not CCT2, loss-of-function in neuron-like SH-SY5Y cells increased intracellular β-amyloid load. These data suggest that CCT2/CCT3 are faithful disease-state indicators and implicate CCT3 in oxidative stress-dependent cellular damage pathways.

## Introduction

Alzheimer’s disease (AD) is broadly recognized as a ‘proteinopathy’, a term coined to mechanistically describe intracellular events when protein production, trafficking, maturation, use, degradation, and secretion become impaired, thus curtailing proteostasis. Accordingly, the main pathological hallmarks of AD are aggregates of β-amyloid (Aβ) and hyperphosphorylated tau, among several other aggregating proteoforms of synaptic, mitochondrial, membrane, and extracellular matrix proteins (Lutz & Peng, 2018). Thus, consensus exists on recognizing the post-translational processing of proteins as a major site of biochemical disruption in neurons in AD.

To become functionally active, proteins ought to fold into unique three-dimensional structures. This process relies on a complex machinery of chaperones that are recruited to an initially nascent polypeptide chain. Cytosolic chaperones and chaperonins accumulate in either the endo-plasmic reticulum or the cytoplasm to hold newly synthesized peptide chains in a state compatible with folding upon release into the cytosol. To do so, eight subunits of chaperonins (CCT1-8) assemble into a double-ringed multi-subunit complex in eukaryotes (that is, the TCP-1 ring complex), which participates in the folding of 10-15% of all newly synthesized proteins (Thulasiraman *et al*., 1999). Besides its cardinal role in proteostasis, TCP-1 activity also prevents the aggregation and unwanted complexing of proteins (for review see; Hartl *et al*., 2011; Gestaut *et al*., 2019). It is not unexpected, therefore, that altered chaperone and chaperonin functions have been associated with brain malformations (Kraft *et al*., 2024), and human brain diseases that are by now classified as ‘proteinopathies’: Huntington’s disease (Tam *et al*., 2006; Darrow *et al*., 2015), Parkinson’s disease (Sot *et al*., 2017), and AD (Slavotinek & Biesecker, 2001; Khabirova *et al*., 2014). Accumulating evidence links specific CCT subunits to a range of processes within the protein quality control system and protein aggregation *per se*. CCT2/5/7 are implicated in autophagy (Pavel *et al*., 2016), with CCT2 identified as an aggrephagy receptor for solid protein aggregates (Ma *et al*., 2022a). In contrast, CCT3/6/7 inhibit protein aggregation (Sot *et al*., 2017; Ben-Maimon *et al*., 2025; Jiang *et al*., 2025).

Transcriptomics studies found altered CCT2-6 mRNA expression (Blalock *et al*., 2004; Ma *et al*., 2022a) and protein concentration (Schuller *et al*., 2001; Xu *et al*., 2019) as abnormalities associated with AD. Nevertheless, neither the neuropathological relevance of CCTs to AD progression at cellular resolution nor their functional interrogation has been performed. Here, we addressed if changes in the expression and subcellular localization of CCTs correlates with either Aβ or tau, or both, in the human brain. For analysis, we selected ATP-dependent CCT2 associated with autophagy and protein aggregate clearance, and CCT3, an inhibitor of protein aggregation. We also asked if impaired CCT availability and function could affect Aβ turnover. We found that neuronal CCT levels and intracellular localization correlated with AD severity, including ectopic expression in astrocytes. CCT3 expression was induced by oxidative stress, which we qualified, based on loss-of-function models, as an adaptive mechanism to limit intracellular Aβ accumulation.

## Materials and methods

### Human brains

*Post-mortem* hippocampi from patients with AD and age-matched controls (without clinical signs of neuropsychiatric disease) were obtained at the London Neurodegenerative Diseases Brain Bank, Institute of Psychiatry, Psychology & Neuroscience, King`s College London. Local ethical committee approval (#231/2001) and informed consent from the next of kin have been obtained to harvest *post-mortem* tissues (**Supporting Table 1**). Neuropathological and histochemical examination, including the classification of Aβ load, and tau pathology, was performed to assess the Alzheimer`s disease neuropathological change. Tissue blocks or entire hippocampi were immersion fixed in 4% paraformaldehyde in 0.1M phosphate buffer (pH 7.4) for 48–72h at 4 °C, followed by long-term storage in 0.1M phosphate buffer (PB) to which 0.1% NaN_3_ had been added. Alternatively, native tissue blocks (2–3 cm^3^) of 1 cm thickness from the opposite hemisphere were stored at −80 °C until processing for biochemical analysis.

### Immunohistochemistry

Protein epitope signature tags (PrESTs) for CCT2 (HPA003197, amino acid 392-519) and CCT3 (HPA006543, amino acid 244-370) were designed using bioinformatic tools with information available on the human genome. Selected PrESTs were recombinantly produced in *E. coli* and used to immunize rabbits. Mono-specific polyclonal antibodies were generated through immunoaffinity purification of the resulting antisera (**Supporting Fig. 1**).

Seven-μm-thick sections were cut on a sliding microtome from paraformaldehyde-fixed paraffin-embedded tissues and mounted on glass slides coated with 3-aminopropyltriethoxysilane. Sections were dewaxed in xylene (10 min), then rehydrated in a descending alcohol gradient (100%, 95%, 70%, and then 50%). Endogenous peroxidase activity was blocked by 1% H_2_O_2_ in methanol (10 min). Antigen retrieval was by 5% urea in 0.1M Tris-HCl (pH 9.5, microwave: 800W, 2 x 6 min) followed by incubation in ice-cold fresh 0.5% NaHBO_4_ in 0.1M PB (pH 7.4).

For immunohistochemistry, slides (or coverslips in cell culture experiments) were incubated in 0.1M PB (pH 7.4) containing 10% normal donkey serum (NDS), 5% BSA, and 0.3% Triton X-100 (all from Sigma) at 22–24 °C for 1h. Subsequently, tissues were exposed to cocktails of primary antibodies (**Supporting Table 2**) in 0.1M PB also containing 1% NDS and 0.2% Triton X-100 at 4 °C overnight. For CCT3/ glial fibrillary acidic protein (GFAP) double-labelling, sections were sequentially stained, first with an anti-CCT3 antibody (rabbit host) and carbocyanine (Cy)3-conjugated donkey anti-rabbit secondary antibody (1:300, 2h; Jackson ImmunoResearch), followed by exposure to biotin-tagged rabbit anti-GFAP antibody overnight. GFAP localization was revealed by using Cy2-conjugated streptavidin (1:500, 2h; Jackson ImmunoResearch). For human brain sections, lipofuscin autofluorescence was blocked by 10% Sudan black B (Merck) diluted in 70% ethanol. Samples were coverslipped by aquamount (DAKO).

### Imaging

Samples were first inspected on a Nikon Eclipse E600 fluorescence microscope (Nikon), equipped with appropriate filter sets for epifluorescence microscopy. Photomicrographs were taken on a Hamamatsu ORCA-ER C4742-80 digital camera (Hamamatsu). Then, confocal laser-scanning microscopy was performed on a Zeiss 710 platform (Zeiss) equipped with appropriate excitation and emission filters for the maximum separation of Cy2, Cy3, and Cy5 signals. Digital images were uniformly optimized for resolution, brightness, and contrast using Photoshop (Adobe Systems) to best reflect visual impressions as observed in the microscope. The intensity of immunoreactivity and signal distribution at the single cell level were measured off-line using ImageJ (NIH) on unprocessed, non-saturated images. For *in vitro* experiments, mean optical density (mean O.D.) for all channels was quantified in randomly selected cells from at least 2 coverslips. Linear intensity plots were generated using the plot profile function.

### *In vitro* H_2_O_2_ challenge

Human U251 glioblastoma-derived cells were plated at a density of 25,000 cells per poly-D-lysine-coated coverslip in 24-well plates in DMEM/F12 + B27 minus antioxidants for 24h. Cells were stimulated with 0, 25, or 50 µM H_2_O_2_ for 24h. Cell cultures were quickly washed in ice cold PBS and fixed in PBS containing 4% paraformaldehyde and stored in PBS containing 0.1% NaN_3_ at 4°C until processing.

### *In vitro* siRNA-mediated loss-of-function experiments

Human SH-SYSY neuroblastoma cells were seeded at a density of 25,000 cells on poly-D-lysine coated coverslips in 24-well plate. Cells were allowed to attach and differentiate for 24h in DMEM/F12 medium (Invitrogen) containing 1 µM retinoic acid (Sigma). Cell cultures were washed with Accell medium (Dharmacon) and incubated in Accell medium containing 1 µM pooled siRNA (Dharmacon, **Supporting Fig. 3**) targeting human CCT2, CCT3 or a mix of CCT2/CCT3 for either 3h or 12h. Control cultures were grown in Accell medium without siRNA for 12h. Cell cultures were quickly washed in ice-cold PBS and fixed in PBS containing 4% paraformaldehyde and stored in PBS containing 0.1% NaN_3_ at 4°C.

### Western blotting

Tissue samples were homogenized in ‘TNE’ buffer containing 1% β-octylglucoside, 5 mM NaF, 100 µM Na_3_VO_4_, 0.5% Triton X-100, and protease inhibitors (‘Complete’, Roche) in a closed-chamber Ultra Turrax^®^ tube drive (IKA). Homogenates were then diluted to a final protein concentration of 2 μg/μl, and denatured. Proteins were resolved by SDS-polyacrylamide gel electrophoreses on 8% gels and transferred onto PVDF membranes (Immobilon-FL, Merck/Millipore). Membranes were probed with primary antibodies (**Supporting Table 2**) followed by exposure to IRDye^®^-800CW and IRDye^®^-680-conjugated secondary antibodies (Li-cor). Proteins of interest were detected and analysed on an Odyssey infrared imager (Li-cor).

### Statistics

Results were analysed using SPSS (SPSS Inc.) and pandas, an open-source data analysis and manipulation tool for Python. Data were expressed as means ± s.e.m. Both the intensity of immunoreactivity and protein expression upon Western blotting were analysed using Welch’s independent-sample *t*-test comparing control *vs*. AD cases at defined stages of the dis-ease. A relationship between GFAP and CCT3 load was determined by linear regression analysis, with the Pearson’s correlation coefficient noted. Data from cell cultures were statistically evaluated using one-way ANOA. A *P* value of < 0.05 was taken as indicative of statistical significance throughout.

## Results

### Study design: specifications and potential limitations

This report was limited to addressing the distribution of chaperonins of the TCP-1 ring complex, particularly CCT2 and CCT3, in the human hippocampus. To do so, affinity-purified antibodies generated within the Human Protein Atlas (HPA) project (*for details see*: **Supporting Fig. 1**) were used. Brain-wide mapping of either target was not performed because broad protein distribution maps after chromogenic detection are available from the HPA consortium, with CCT2 (https://www.proteinatlas.org/ENSG00000166226-CCT2) and CCT3 (https://www.proteinatlas.org/ENSG00000163468-CCT3) classified as proteins with low tissue and brain region specificity as chaperonin functions are essential for all cells. Instead, in-depth cellular and subcellular analyses were emphasized in relation to the progressive pathobiology of AD. CERAD-based terminology was used throughout, with designations as ‘normal’ (control), ‘possible AD’ (Braak stage I-II), ‘probable AD’ (Braak stage III-IV), and ‘definite AD’ (Braak stage V-VI; **Supporting Table 1**).

### Distribution of CCT2 in the hippocampus of control and AD subjects

In control subjects (*see also* **Supporting Table 1,2**), CCT2-like immunoreactivity dominated in neurons in the hippocampal formation, including those positioned in the granular and polymorph layers of the dentate gyrus (**Fig. 1a**), and pyramidal cells of the CA subfields (CA1, **Fig. 1b**). CCT2-like immunoreactivity was restricted to the soma and primary apical and basal dendrites of neurons, with no labelling seen in either nuclei or axons/terminal boutons. CCT2 colocalized with neither integrin subunit alpha M (ITGAM, CD11b) nor glial-fibrillary acidic protein (GFAP), suggesting predominant CCT2 expression in neurons.

**Figure 1.**
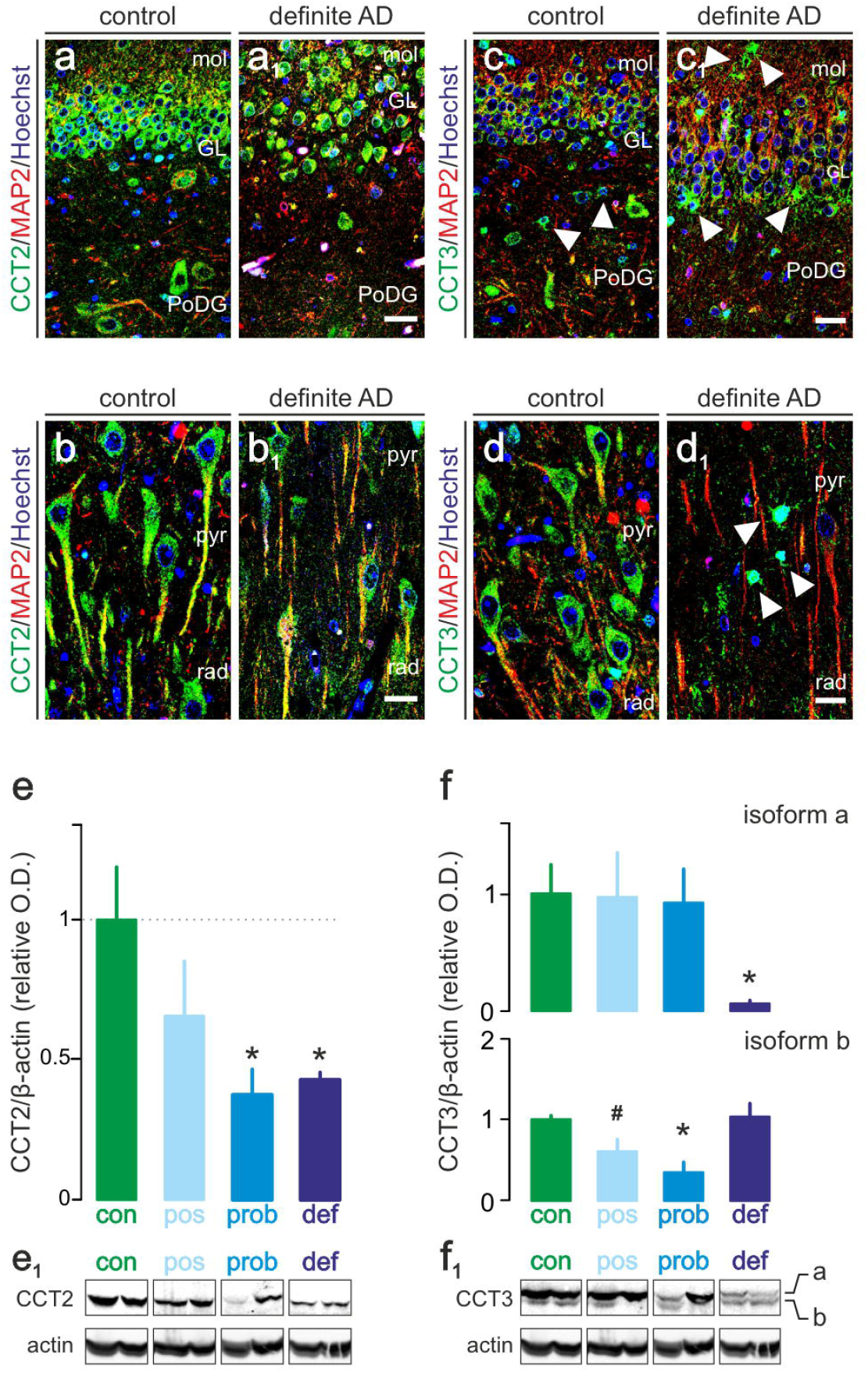
CCT2 and CCT3 in the human hippocampus at stratified AD stages. (**a**) In the dentate gyrus, CCT2 accumulated in granule cells, pyramidal cells of the CA4 sub-field, and hilar neurons. (**a**_**1**_) CCT2-like immunoreactivity was reduced as the granule cell layer became disorganized in definite AD. (**b**) CCT2 in CA1 pyramidal neurons, particularly in their somata and proximal dendrites. (**b**_**1**_) Here, CCT2-like immunoreactivity was also reduced in definite AD. (**c**) In AD, an increased number of tangle-bearing neurons and extracellular plaques (*arrowheads*) were associated with reduced CCT3-like immunoreactivity in neurons in, e.g., the polymorph layer of the dentate gyrus (PoDG; **c**_**1**_). In definite AD (**d**_**1**_ *vs*. **d**), perisomatic CCT3-like immunoreactivity was absent in those CA1 pyramidal neurons that were proximal to extracellular plaques (*arrowheads*). (**e,f**) Western blotting revealed gradually reduction in both CCT2 and CCT3 protein levels as a factor of disease progression in the human hippocampus. For CCT3, two isoforms (‘a’ (75 kDa) ‘b’ (65 kDa), corresponding to possible splice variants) were quantified. β-Actin was used as a house-keeping standard. Representative Western blots were shown (e_1_,f_1_) with a total of *n* = 21 cases quantified. ^#^*p* < 0.1, \**p* < 0.05, ** *p* <0.001. *Abbreviations*: cont, control; def, definite AD; GL, granule cell layer; mol, molecular layer; pos, possible AD; prob, probable AD; pyr, pyramidal layer; rad, radial layer. *Scale bars* = 40 µm (a_1_,c_1_), 15 µm (b_1_,d_1_).

Perisomatic CCT2-like immunoreactivity remained unchanged in the granular cell layer of the dentate gyrus in both ‘probable AD’ and ‘definite AD’ with clear histopathological indices of disease progression in terms of both hyperphosphorylated tau and senile plaque formation intra- and extracellularly, respectively (**Fig. 1a_1_**). Nevertheless, CCT2-like immunoreactivity was reduced in the dendrites of pyramidal cells in the CA1 subfield (**Fig. 1b_1_**). A decrease in CCT2-like immunoreactivity was prominent in neurons harbouring hyperphosphorylted tau, particularly in ‘definite AD’ subjects (**Supporting Fig. 2a-c**_**2**_**’’**).

Subsequent Western blotting validated our histochemical results for which ‘normal’ (*n* = 4), ‘possible’ (*n* = 5), ‘probable’ (*n* = 6) and ‘definite’ AD cases (*n* = 6) were analysed in parallel. Western blotting identified a single CCT2-immunoreactive(^+^) band of ~56 kDa in human tissue homogenates, which corresponds well to the predicted size of this protein (535 amino acids; 57.488 kDa; UniProt). As **Fig. 1e,e_1_** show, CCT2 tissue levels gradually decreased as a factor of the severity of AD. This reduction reached statistical significance at both the ‘probable’ (*t* = 3.627, *p* = 0.014) and ‘definite’ (*t* = 3.829, *p* = 0.028) AD groups *vs*. ‘normal’ subjects. In sum, CCT2 protein levels are negatively affected by AD, particularly in neurons that show indices of cellular damage.

### Distribution of CCT3 in the hippocampus of control *vs*. AD subjects

Like CCT2, CCT3 was found, chiefly, in granule cells of the dentate gyrus (**Fig. 1c**), and the somata and dendrites of pyramidal cells of the CA subfields (CA1, **Fig. 1d**). Nuclei remained unstained throughout. In addition, CCT3-like immunoreactivity was seen in a subset of polymorph cells, with morphologies reminiscent of astrocytes. CCT3 did not co-localize with IT-GAM, excluding expression by microglia (*data not shown*).

In ‘definite AD’, both perisomatic and dendritic CCT3-like immunoreactivity became reduced in pyramidal cells in the CA1 subfield (**Fig. 1d_1_; Supporting Fig. 2e-f**), with dentate granule cells being less affected. Notably, star-shaped cells apposing tau-laden proximal dendrites in the CA1 subfield were frequently encountered in ‘definite AD’, too (**Fig. 1d_1_**). Dual labelling with GFAP confirmed CCT3-like immunoreactivity in astrocytes (*see below* and **Fig. 4**).

Next, we applied Western blotting to test tissue CCT3 levels across stratified subjects as above. Human CCT3 has 6 known splice variants (60, 57, 56, 30, 17 and 17 kDa). Two CCT3^+^ bands were found by Western blotting (termed ‘a’ (75 kDa) an ‘b’ (65 kDa) isoforms). CCT ‘a’ isoform levels were unaltered in ‘possible’ and ‘probable’ AD but significantly reduced in ‘definite’ AD subjects (*t* = 4.108; *p* = 0.004; **Fig. 1f,f_1_**). In contrast, the 65 kDa product of CCT3 (‘b’) showed a gradual decrease, reaching significance at ‘probable’ (*t* = 3.337; *p* = 0.011) and ‘definite’ stages of AD (*t* = 3.753; *p* = 0.006; **Fig. 1f,f_1_**).

### Subcellular distribution of CCT2 and CCT3 in Aβ-bearing neurons

Neurodegeneration changes the cellular composition of the hippocampus. We performed high-resolution confocal microscopy along the somatodendritic axis of CA1 pyramidal cells (**Fig. 2a,b**) exhibiting a gradient of intracellular Aβ inclusions to determine if CCT2 and/or CCT3 levels and subcellular distribution were affected. Therefore, pyramidal cells were co-labelled for either CCT2 or CCT3 and Aβ in a case (C06) with early pathological indices of AD. Planar, thin optical sections (<0.4 µm optical thickness) were captured by confocal microscopy. For CCT2, immunolabelling appeared unperturbed and homogeneous at the light microscopy level in pyramidal cell bodies with a handful (<5) of Aβ(17-25)^+^ puncta (**Fig. 2c**). In contrast, increasing Aβ load was associated with the focal exclusion of CCT2 from the cytosol (**Fig. 2c,d**). The intracellular redistribution of CCT3 was reminiscent of CCT2 (**Fig. 2e,f**). These data show that *de novo* Aβ synthesis and intracellular accumulation are associated with the exclusion of chaperonin subunits from mainly perinuclear cellular compartments in AD.

**Figure 2.**
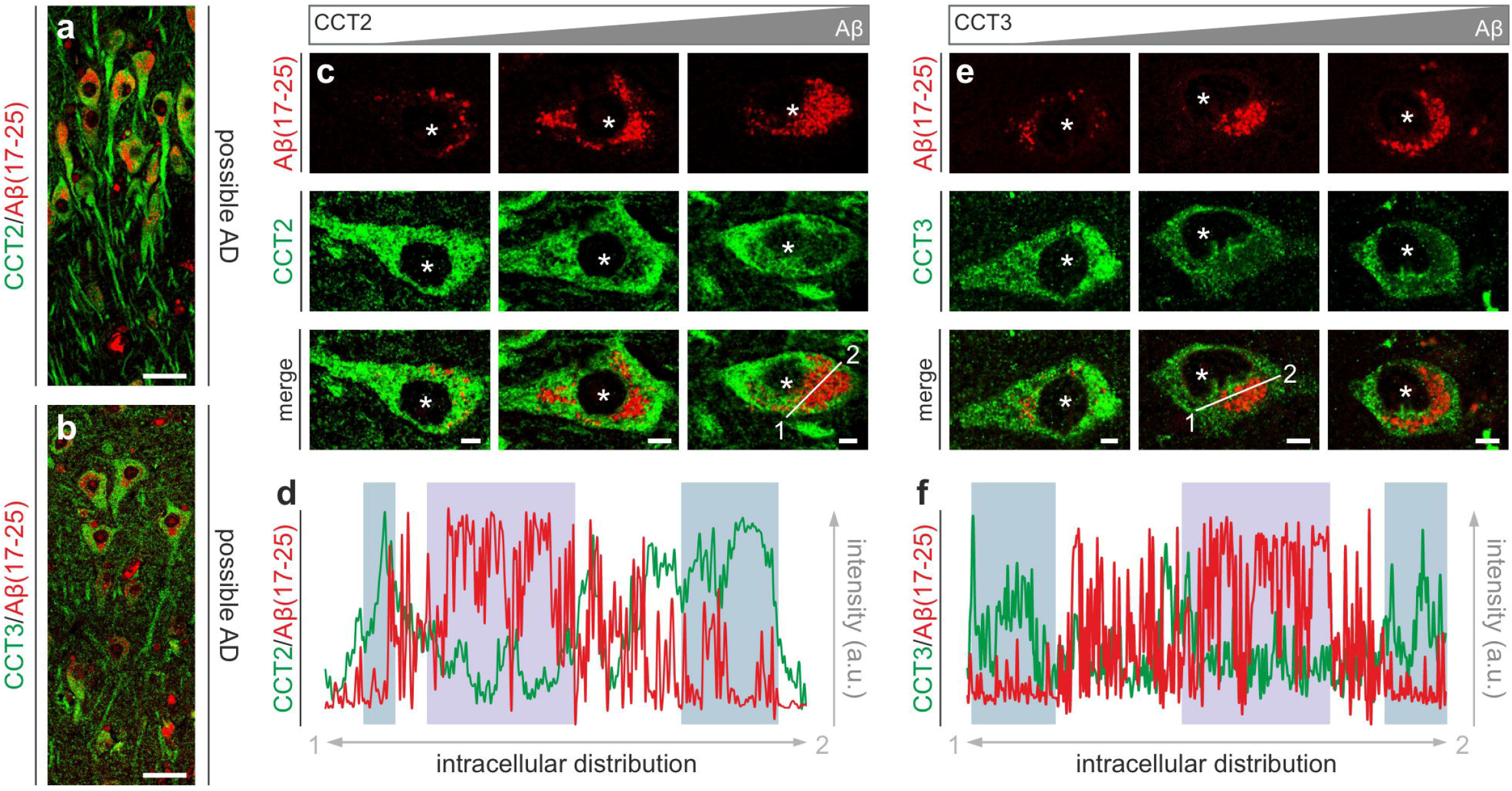
Subcellular CCT2 and CCT3 distribution in relation to intracellular Aβ. (**a,b**) Intracellular Aβ accumulation in CA1 pyramidal neurons was found as early as in possible AD. (**c**) CCT2-like immunoreactivity was mutually exclusive with Aβ intracellularly; as was confirmed by plotting fluorescence signal intensity in affected neurons (along the line between 1,2 (example); **d**). (**e**) Similarly, CCT3-like immunoreactivity was excluded from Aβ-laden sub-cellular areas (**f**). Asterisks denote the location of nuclei. *Scale bars* = 15 µm (a,b), 2 µm (e,f).

### Subcellular distribution of CCT2 and CCT3 in neurons with neurofibrillary tangles

We used an identical approach to analyse the relationship between subcellular CCT2/CCT3 and hyperphosphorylated tau distribution. **Fig. 3a,b** show the complementary intracellular localization of CCT2 and paired-helical filaments in hippocampal neurons. Both CCT2 and CCT3-like immunoreactivities were homogeneous in the cytosol of tangle-negative neurons. Once perinuclear tangles formed, they juxtaposed with CCT2/CCT3-like immunoreactivities, which were reduced in both density and intensity. Indeed, pervasive tau pathology led to the absence of both CCT2 and CCT3 in pyramidal neurons (**Fig. 3c,d**).

Next, we used a population approach to quantify CCT2-like immunoreactivity in pyramidal neurons with or without hyperphosphorylated tau immunoreactivity (**Fig. 3e**) using within-sample comparisons. CCT2-like immunoreactivity in the perikarya of neurons with tau inclusions was significantly reduced (*t* = 3.136; *p* = 0.007) even prior to the appearance of classical neurofibrillary tangles (**Fig. 3f**). In sum, these data reinforce disrupted protein folding in tangle-bearing neurons in AD.

**Figure 3.**
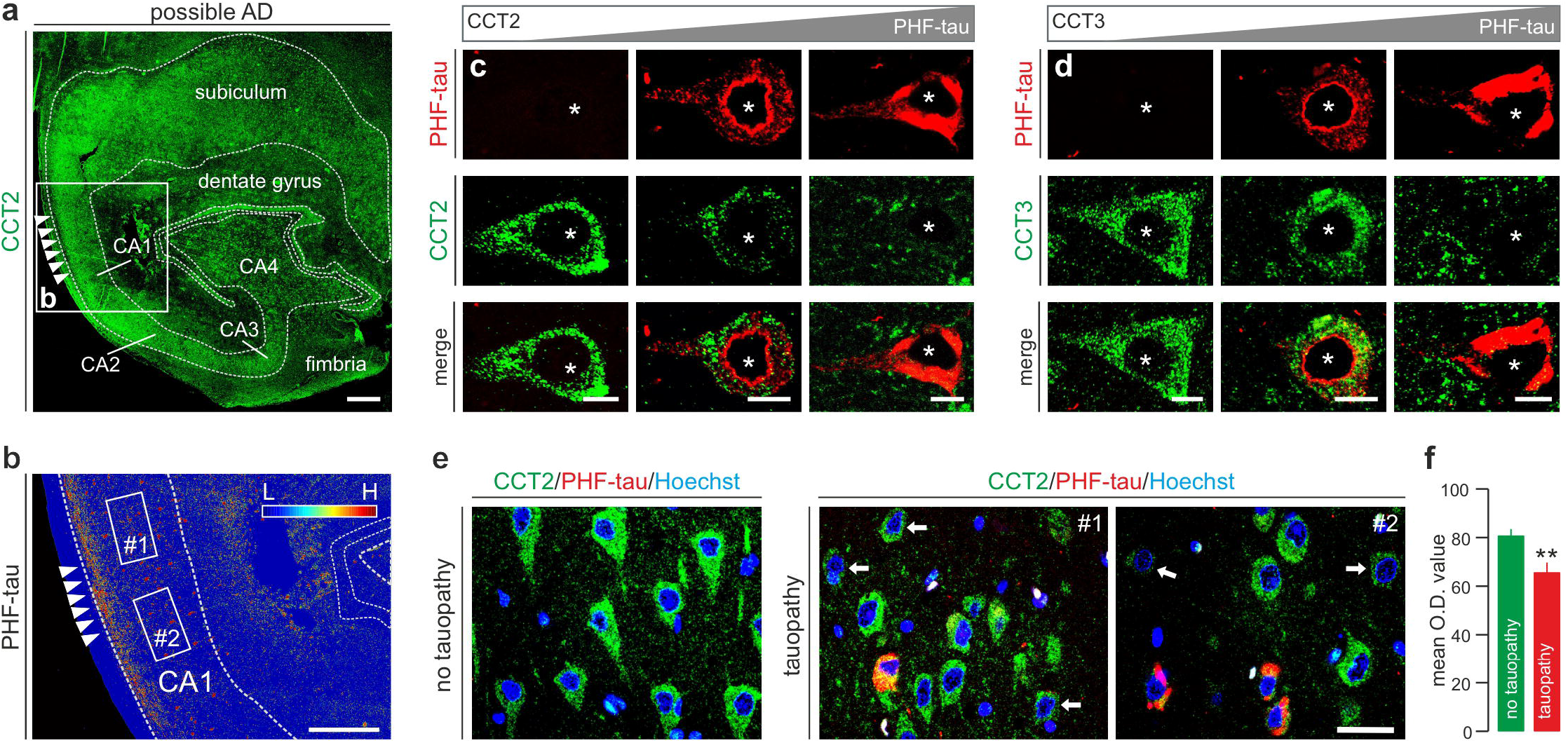
CCT2 and CCT3 immunoreactivity in relation to tau pathology. (**a**) Overview of the hippocampal formation and the parahippocampal gyrus immunostained for CCT2 from a subject with possible AD. (**b**) Heat-map of tau immunoreactivity color-coded from low (blue) to high (red) abundance. *Arrowheads* point to a tau-bearing locus. #1 and #2 are shown at higher magnification in panel (e). (**c,d**) The accumulation of hyperphosphorylated tau commenced in the perinuclear cytoplasm, with a gradual spread towards dendrites. Both CCT2-like and CCT3-like immunoreactivities were reduced subcellularly where tau accumulation occurred (*arrows*). Asterisks denote the positions of nuclei. (**e**) Comparison of CCT2-like immunoreactivity in cell ensembles in hippocampal CA1 in control *vs*. possible AD. (**f**) CCT2-like immunoreactivity was significantly reduced (\**p* <. 0.05). *Scale bars* = 100 µm (a), 50 µm (b), 17 µm (e), 2 µm (c,d).

### CCT3 in astrocytes

Besides neurons, CCT3 was also expressed in protoplasmic astrocytes (**Fig. 4a,d**). The population of CCT3^+^ astrocytes was sparse throughout the hippocampal formation even in some ‘normal’ cases. In turn, CCT3^+^ astrocytes where frequently positioned proximal to tau-bearing neurons in AD (**Fig. 4e**). Therefore, we asked if a relationship between GFAP^+^ (‘reactive’) astrocytes and CCT3 load could be established through coincident detection of CCT3 (**Fig. 1g,h**) and GFAP (**Fig. 4b**) by Western blotting and subsequent correlation of their tissue levels. Both full-length (50 kDa) and truncated GFAP (40 kDa) were detected (**Fig. 4b**). Correlation analysis revealed a close relationship between 40 kDa GFAP degradation products and *post-mortem* delay (R^2^ = 0.564; *p* = 0.008). Therefore, full-length (50 kDa) GFAP was used throughout. GFAP tissue content did not vary significantly across AD stages in our subject cohort (**Fig. 4b**). When assessing a possible correlation between GFAP and CCT3 ‘a’, which is the dominant isoform in human brain (**Fig. 1h**), no relationship was seen in either ‘normal’ (R^2^ = −0.105; *p* = 0.895) or ‘definite’ AD subjects (R^2^ = −0.050; *p* = 0.925) because CCT3 load was either preserved (‘normal’) or invariably reduced (‘definite’). In contrast, a significant positive correlation existed in cases with either possible (R^2^ = 0.977; *p* = 0.004) or ‘probable’ AD (R^2^ = 0.816; *p* = 0.048; **Fig. 4c**). These data suggest adaptive astroglial CCT3 expression in AD.

**Figure 4.**
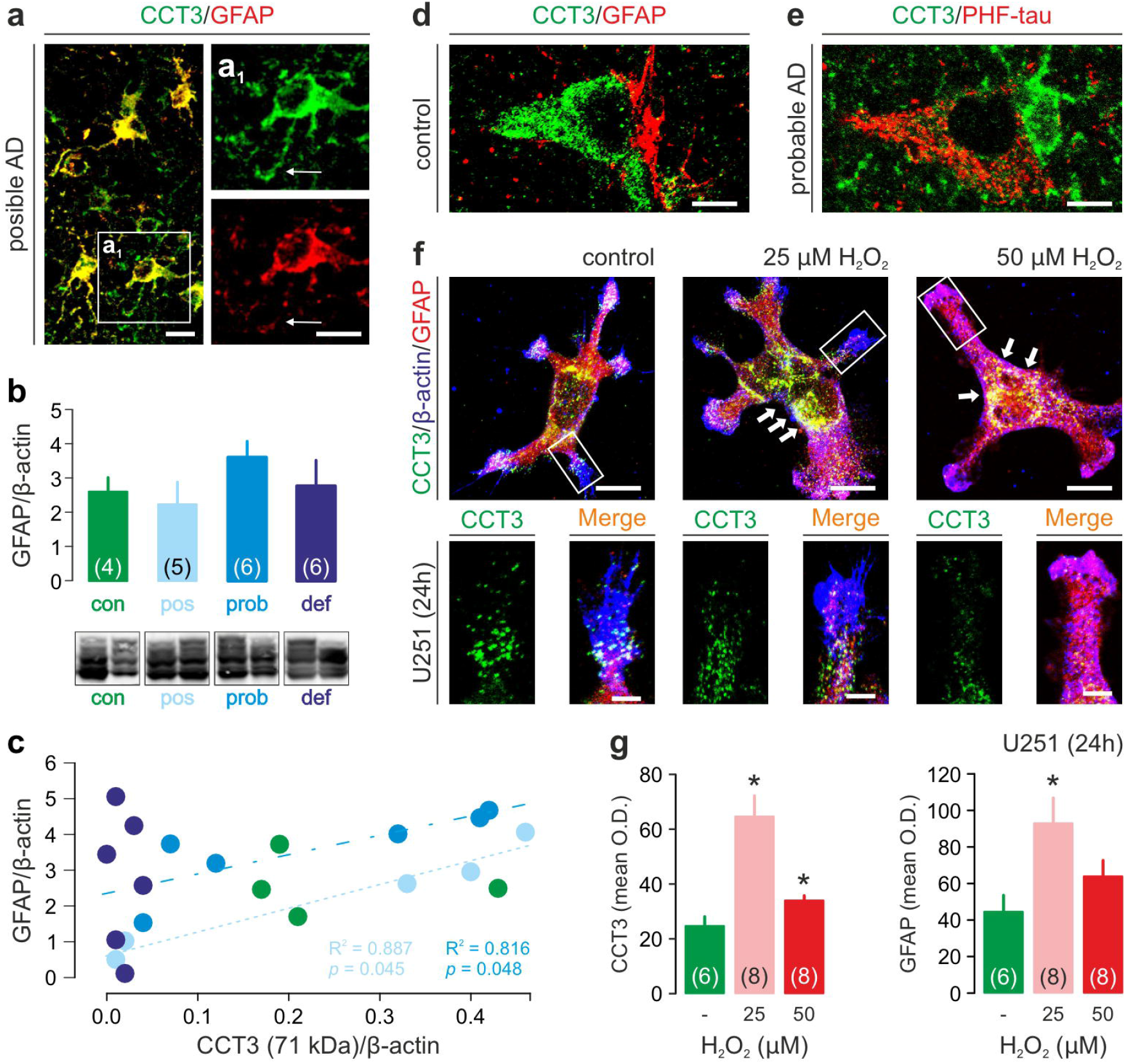
CCT3 expression in astrocytes and its sensitivity to H_2_O_2_. (**a,a**_**1**_) Besides neurons, CCT3-like immunoreactivity was also seen in astrocytes positive for glial fibrillary acidic protein (GFAP). Note the differential localization of CCT3 and GFAP in astrocyte processes (*arrows*). (**b**) GFAP content was unchanged in AD. Bracketed numbers indicate case numbers per group. (**c**) Positive correlation between GFAP and CCT3-like immunoreactivities in subsets of AD cases. The Pearson correlation co-efficient was provided for ‘possible’ and ‘probable’ AD cohorts. (**d**) CCT3^+^ neuron apposing a CCT3^-^ astrocyte. (**e**) CCT3^+^ profuse cell morphology in the proximity of a tangle-bearing dystrophic neuron. (**f**) U251 astrocytoma cells in control conditions expressed CCT3 *in vitro*, including their processes and lamellipodia (*insets*). When challenged with either 25 µM or 50 µM H_2_O_2_, CCT3-like immunoreactivity was reduced in astroglioma processes. (**g**) At the same time, perisomatic CCT3 (*arrows*) and GFAP immunoreactivities significantly increased, as was also shown quantitatively (bottom row; \**p* < 0.05). Numbers in brackets show the number of independent cellular reconstructions. *Scale bars* = 10 µm (a,a_1_), 7 µm (f, *overviews*) 2 µm (d,e, f/*insets*).

### Cellular redistribution of CCT3 in U251 cells challenged by H_**2**_**O**_**2**_

We asked if CCT3 expression is an inducible feature of reactive astrocyte-like cells in cellular models that recapitulate oxidative stress-induced cellular damage. To this end, astrocyte-like U251 cells were used because they retain, among others, GFAP expression *in vitro* (Nishiyama *et al*., 1989). To mimic pathology-relevant oxidative stress, U251 cells were exposed to either moderate (25 µM) or elevated (50 µM) H_2_O_2_ concentrations that did not induce apoptosis (Liu *et al*., 2015). Under control conditions, CCT3 immunoreactivity in U251 cells was distributed in the cytosol in a punctate fashion, including processes and lamellipodia (**Fig. 4f**). When exposed to either 25 µM (*t* = 4.803, *p* = 0.001) or 50 µM (*t* = 2.416, *p* = 0.044) H_2_O_2_, cellular CCT3 content was increased relative to controls (**Fig. 4f,g**). Simultaneously, GFAP immunoreactivity also increased, particularly upon exposure to 25 µM H_2_O_2_ (*t* = 2.924, *p* = 0.013, **Fig. 4g**). GFAP immunoreactivity remained evenly distributed, including processes and lamellipodia. In contrast, CCT3 accumulated focally in the perinuclear cytosol while being dose-dependently excluded from processes (**Fig. 4f**). These data suggest that H_2_O_2_-induced oxidative stress increases CCT3 expression and cellular availability, at least, in U251 cells.

### CCT3 gene silencing increases Aβ load in SH-SY5Y cells

Because of the mutually exclusive intracellular localization of Aβ(17-24) and CCT2/CCT3 in hippocampal neurons of AD subjects, and the notion that Aβ misprocessing is due to faulty protein folding, we asked if reducing CCT2 and/or CCT3 levels affected Aβ accumulation *in vitro*. Here, we used SH-SY5Y human neuroblastoma cells, which can produce full-length Aβ when experimentally challenged (Zheng *et al*., 2013). Under control conditions, Aβ(17-24) immuno-reactivity was sporadically seen in the processes of differentiated SH-SY5Y cells (*data not shown*). Upon H_2_O_2_ exposure (12h, **Figure 5a,a_1_**), the amount of Aβ(17-24) immunoreactivity was moderately elevated, mainly in the fine processes of SH-SY5Y cells. Treatment with CCT2 siRNA (**Figure 5b,b_1,_e**) did not affect Aβ(17-24) immunoreactivity in either processes or perikarya at either time point (3h: *t* = 0.337, *p* = 0.740; 12h: *t* = 0.667, *p* = 0.516). In contrast, treatment with CCT3 siRNA significantly increased cellular Aβ(17-24) immunoreactivity after both 3h (*t* = 2.870, *p* = 0.011) and 12h (*t* = 3.213, *p* = 0.007) of H_2_O_2_ exposure (**Figure 5c,c_1_,e**), as compared to H_2_O_2_-only controls at 12h. No apparent additive effect was recorded when knocking down CCT2 and CCT3 simultaneously (**Figure 5d,d_1_,e**; 3h: *t* = 2.131, *p* = 0.051; 12h: *t* = 4.7073, *p* = 0.0003). These data suggest that reducing CCT3 levels in conjunction with oxidative stress increases Aβ(17-24) production.

**Figure 5.**
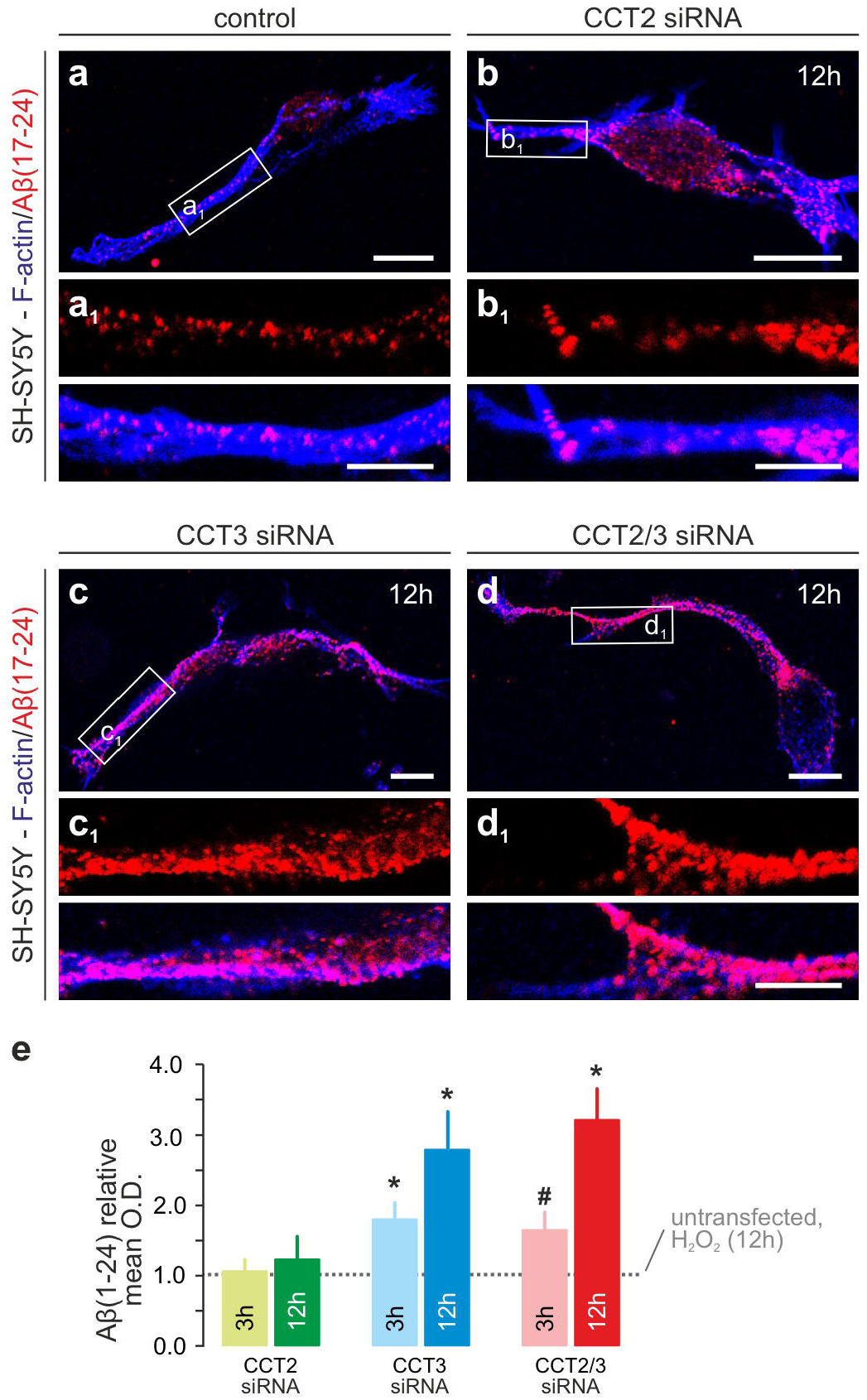
RNAi-mediated CCT2/3 silencing increases intracellular Aβ load. (**a-d**_**1**_) Cellular overviews and high-power photomicrographs of select processes. Open rectangles denote the locations of the insets in (a_1_-d_1_). (**e**) Quantification of intracellular βA immunoreactivity. Note that CCT3 gene silencing significantly increased Aβ(1-24) intracellularly, while manipulating CCT2 expression was ineffective. *Scale bars* = 6 µm (a-d), 1.5 µm (a_1_-d_1_).

## Discussion

Here, we used an antibody-based approach to determine the expression and cellular distribution of CCT2 and CCT3 chaperonin subunits in the human hippocampus, and their changes in AD. Even if earlier transcriptome data suggested the downregulation of both CCT2 and CCT3 in AD, protein-level information on either target remained unknown. By generating affinity-purified antibodies against both CCT2 and CCT3, we first described pyramidal and granule cells as their primary cellular foci of expression in the human hippocampal formation, localized both targets within the somatodendritic axis of neurons, and confirmed their reduced levels in AD. The mutually exclusivity of Aβ, tau *vs*. CCT2/CCT3 in the perisomatic compartment of injured neurons reinforced the disruption of protein processing pathways in AD, qualifying the disease as a ‘proteinopathy’. Moreover, the intra- and subcellular redistribution of CCT2 and CCT3 co-incident with the first incidences of Aβ and/or tau inclusions in neurons support that the break-down of the protein synthesis and maturation/folding machinery substantially contributes to neuronal disfunction. Overall, these findings, together with earlier reports (Blalock *et al*., 2004; Xu *et al*., 2019; Ma *et al*., 2022a; Jiang *et al*., 2025), position CCT subunits as markers to stratify AD at the tissue, regional, cellular, and subcellular levels by faithfully reporting pathological cell states.

Protein mapping and functional studies described both cell-type and cell-state-dependent roles for subunits of the TCP-1 ring complex, including canonical (when the subunits work synergistically) and non-canonical (that is, individual, complex-independent) functions. Unexpectedly, non-canonical functions are more often associated with pathological conditions, particularly because an ‘equivalent loss’ of all TCP-1 complex subunits is rarely reported. For example, retention of CCT3 and CCT7, but not CCT8, can inhibit tau aggregation (Ben-Maimon *et al*., 2025). CCT3 and CCT6, but not other subunits, have affinity to bind α-synuclein (Sot *et al*., 2017). CCT2 and CCT6 but not others modulate the lysosome-dependent clearance of huntingtin (Ma *et al*., 2022b). Our results agree with subunit-specific vulnerabilities in disease because CCT3 but not CCT2 affected intracellular Aβ accumulation under oxidate stress. Cumulatively, these findings indicate an evolutionary gain-of-function for chaperonin subunits to maintain proteostasis and enhancing the survival of post-mitotic cells.

‘Reactive’ astrocytes, classified as GFAP^+^ with hypertrophic somata and processes, are a pathological feature of AD. As AD progresses, astrocytes continuously change their gene expression profiles in an adaptive response to match their local environments (Serrano-Pozo *et al*., 2024). Neuroinflammation (Liddelow *et al*., 2017), and direct damage response pathways to reactive oxygen species (Aksenov *et al*., 2000; Zheng *et al*., 2024) dominate amongst the adaptive features of astrocytes. Increased expression of CCT3 in hippocampal astrocytes, as shown here, could be a consequence of oxidative damage to proteins, a cellular repair response to either clear misfolded proteins or prevent protein aggregation, or both. Our studies *in vitro* support this hypothesis by linking CCT3 expression to oxidative stress.

Neurodegenerative disorders take decades to develop because of the gradual and looped amplification and propagation of pathological processes. Our gene silencing experiments suggest that disrupting CCT3 availability increases intracellular Aβ accumulation. In turn, intracellular aggregates drive a loss of CCT availability, and likely also function, thus self-perpetuating protein aggregation. The role of heat shock proteins (Hsp72, Hsp90, chaperones) in neurodegenerative disorders is well known. Therapies targeting heat shock proteins are currently in development, with some in clinical trials (f*or review see*, Campanella *et al*., 2018). In cancer research, CCT subunits (including CCT2 and CCT3) are recognized as causal to cancer cell survival in harsh tumour environments. Blocking CCT functions reduces tumour cell viability and invasiveness (Cox *et al*., 2022). Therefore, retaining CCT functions in the context of neurodegenerative disorders could be appealing to enhance cell survival by limiting abnormal protein folding and toxicity.

## Funding statement

This work was supported by the Austrian Science Fund (10.55776/COE16, T.Ha.), the Swedish Research Council (2023-02656, J.M.; 2023-03058, T.Ha.), the Swedish Brain Foundation (Hjärnfonden, FO2022-0300, T.Ha.), AlzheimerFonden (AF-1033216, J.M.), the Novo Nordisk Foundation (NNF23OC0084476, T.Ha.), and the European Research Council (FOODFORLIFE, ERC-2020-AdG-101021016; to T.Ha.).

Author contributions

J.M., T.Ho., and T.Ha. designed experiments and conceptualized data. T.Ho. contributed human tissues. J.M. performed all experiments and data analysis. J.M. and T.Ha. procured funding. J.M. and T.Ha. drafted the manuscript with input from T.Ho.

## Legends to Supporting Figures and Tables

**Supporting Figure 1. Antibody generation and specificity**

(**a**) Within the Human Protein Atlas (HPA) project, rabbits were immunized against the amino acid sequences highlighted within CCT2 (2x) and CCT3. (**b**) The specificity of anti-CCT2 antibodies (note their HPA database designations) were first tested in protein Epitope Tag (PrEST) screens. (**c**) Subsequently, antibody specificity was substantiated on Western blots from both cell lines and *post-mortem* hippocampal homogenates.

**Supporting Fig. 2. CCT2 and CCT3 immunoreactivity in control and AD subjects**

Data were presented from hippocampal CA1 with increasing tau load. Note that two cases (designated as #1, #2) per group were included for CCT2 because of the breadth of tau load. Regardless, tau immunoreactivity appeared mutually exclusive for both CCT2 and CCT3 (*arrows*). *Scale bars* = 70 µm (a-c_1_, d-f), 15 µm (a_2_-c_2_), 10 µm (a_2_”-c_2_”).

**Supporting Figure 3. RNAi-mediated gene silencing**

(**a,b**) Mixtures of four siRNAs (CCT2-1—CCT2-4 and CCT3-1—CCT3-4) were pooled and used in each experiment. The sequence specificity of siRNAs is shown. (**a**_**1**_,**b**_**1**_) Detailed information on the target sequences, nucleotide positions, and exon allocation were provided. (**a**_**2**_,**b**_**2**_) Successful silencing at increasing siRNA concentrations was shown by Western blotting 72 h post-transfection. β-Actin served as internal standard.

**Supporting Table 1. Demographic and *post-mortem* pathological information on the subject cohort used in the present study**.

Cases were anonymized throughout. *Abbreviations*: BW, brain wet weight; IH, immunohistochemistry; n.a., not available; PMD, post-mortem delay; WB, Western blotting; y, year.

**Supporting Table 2. Immunoreagents used in the present study**

^1^Paraffin sections from human brain samples. ^2^*In vitro* experiments using U251 and SH-SY5Y cells. *Abbreviations*: HPA, Human Protein Atlas; IHC, immunohistochemistry; n.a., not applicable; WB, Western blotting.

