## Supplementary figures and images for "Stratifying cellular injury in Alzheimer’s disease by chaperonin containing TCP1 subunits 2 and 3"

### Figure S1

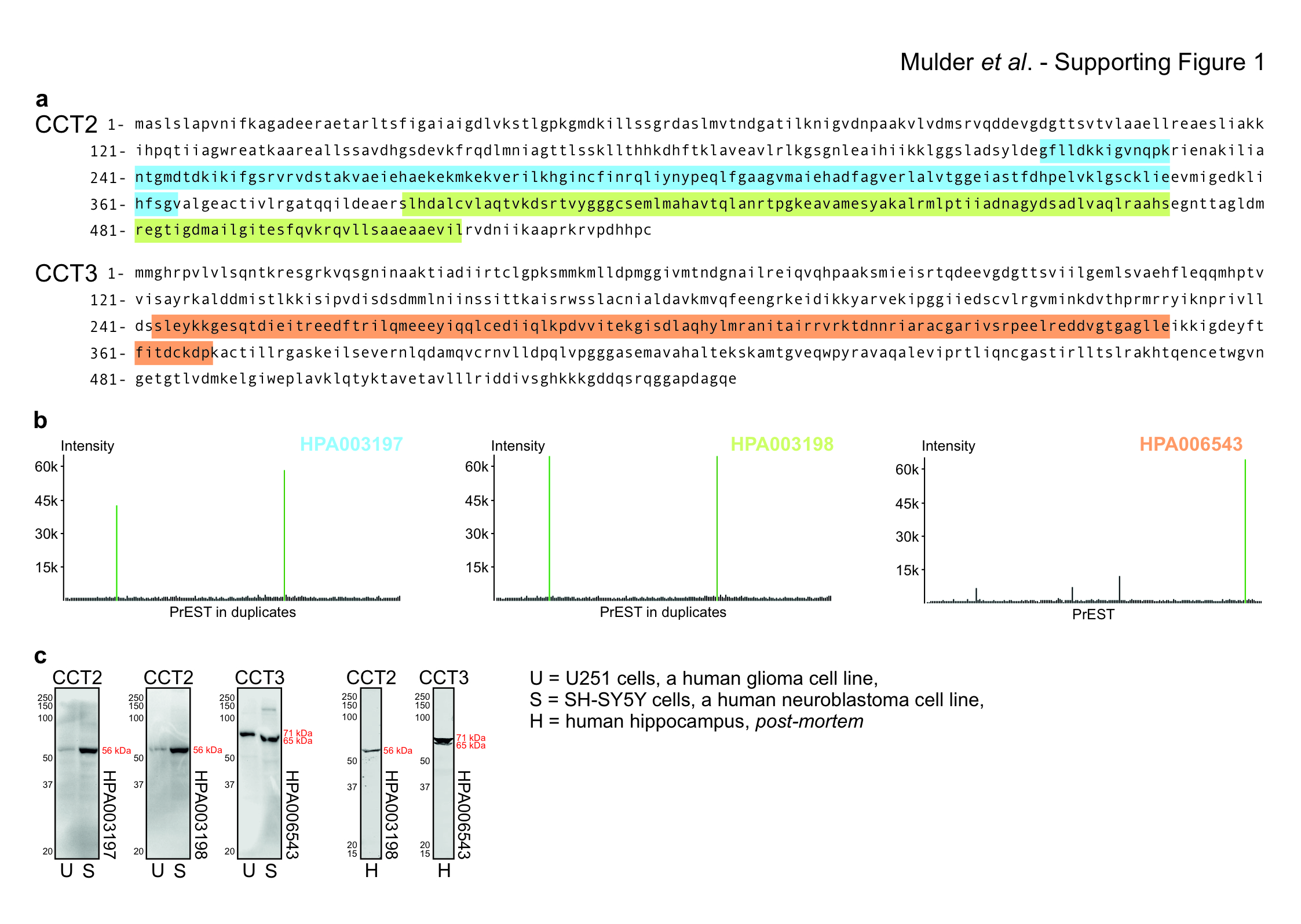

### Figure S2

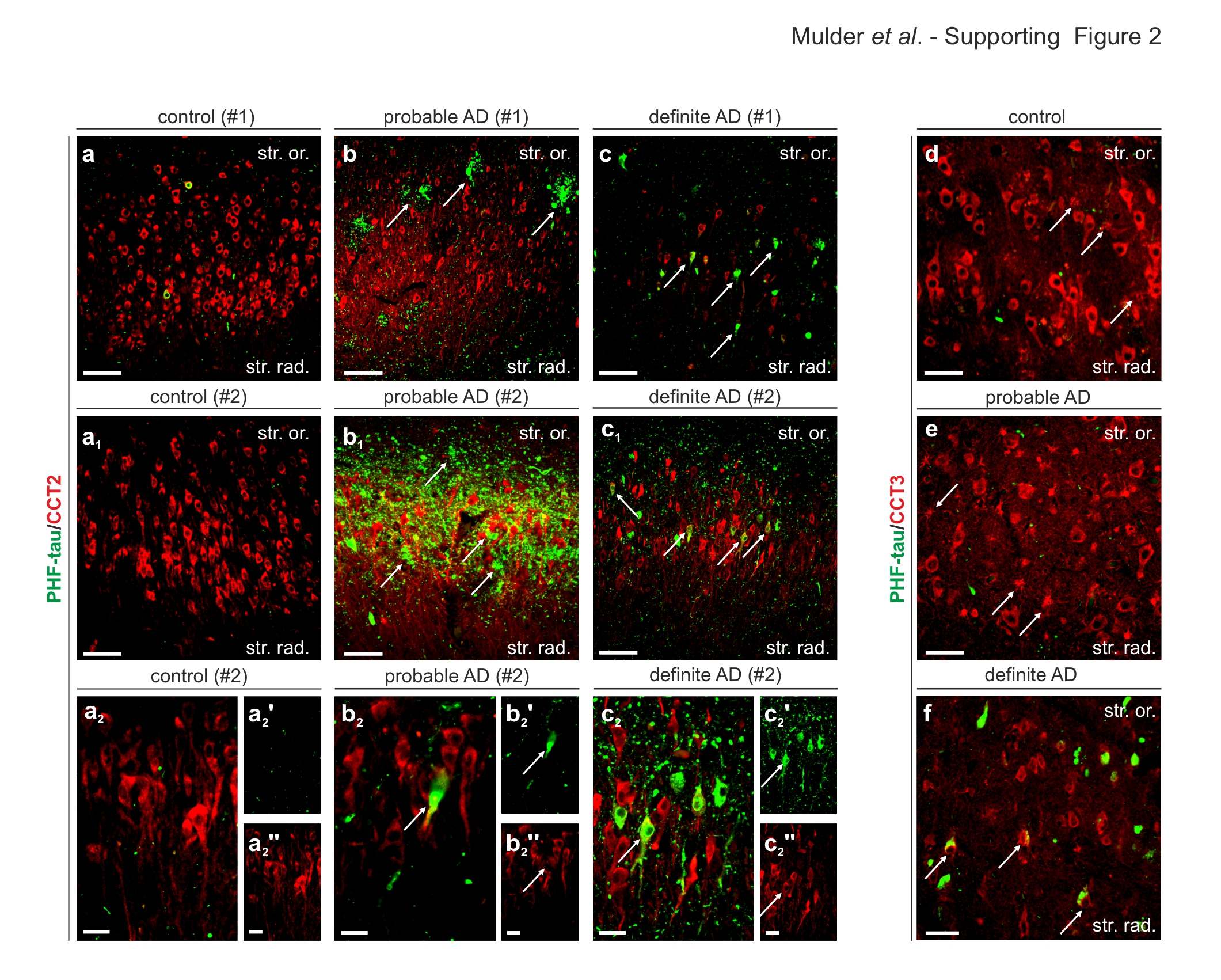

### Figure S3

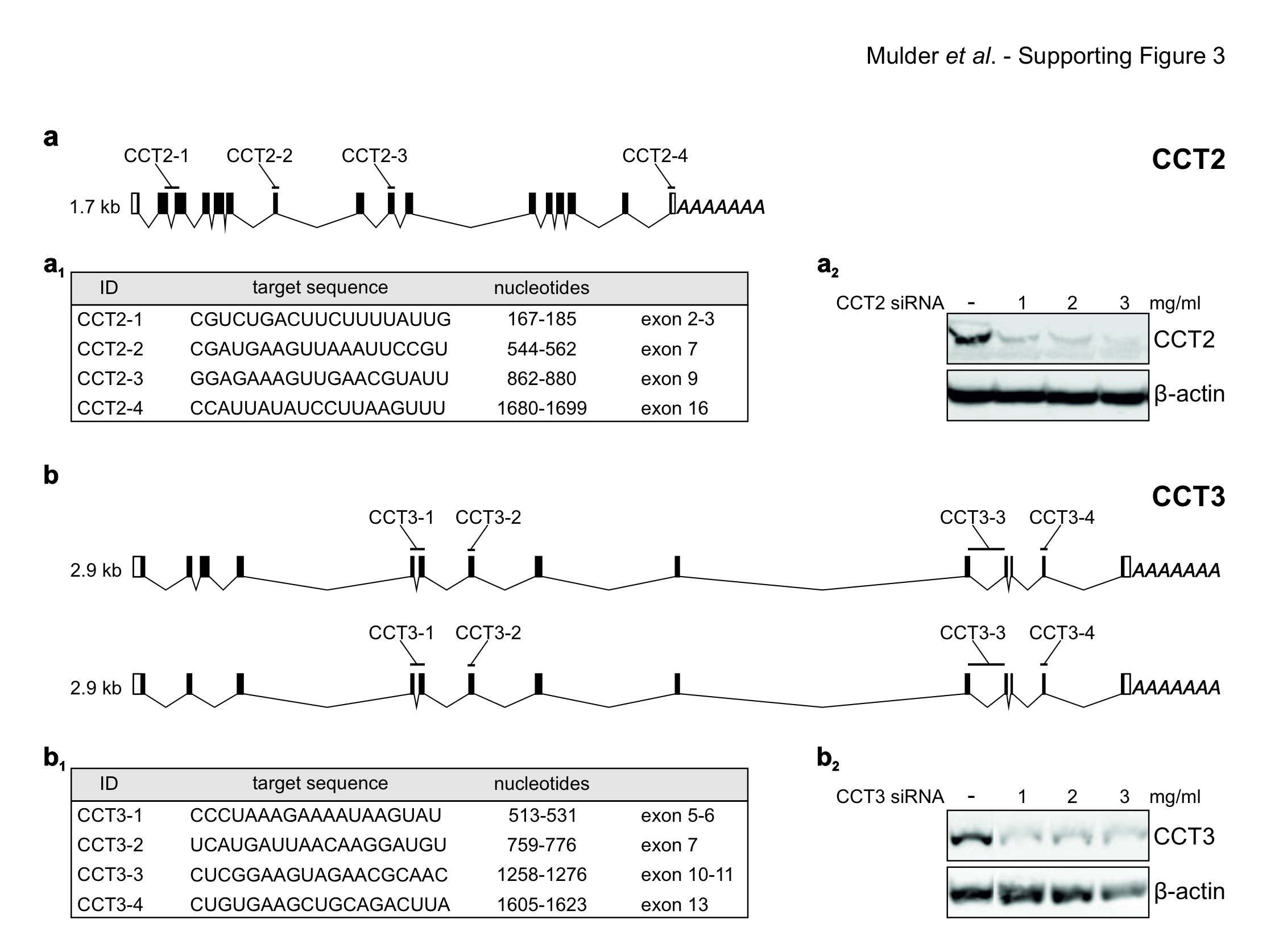

### Table S1

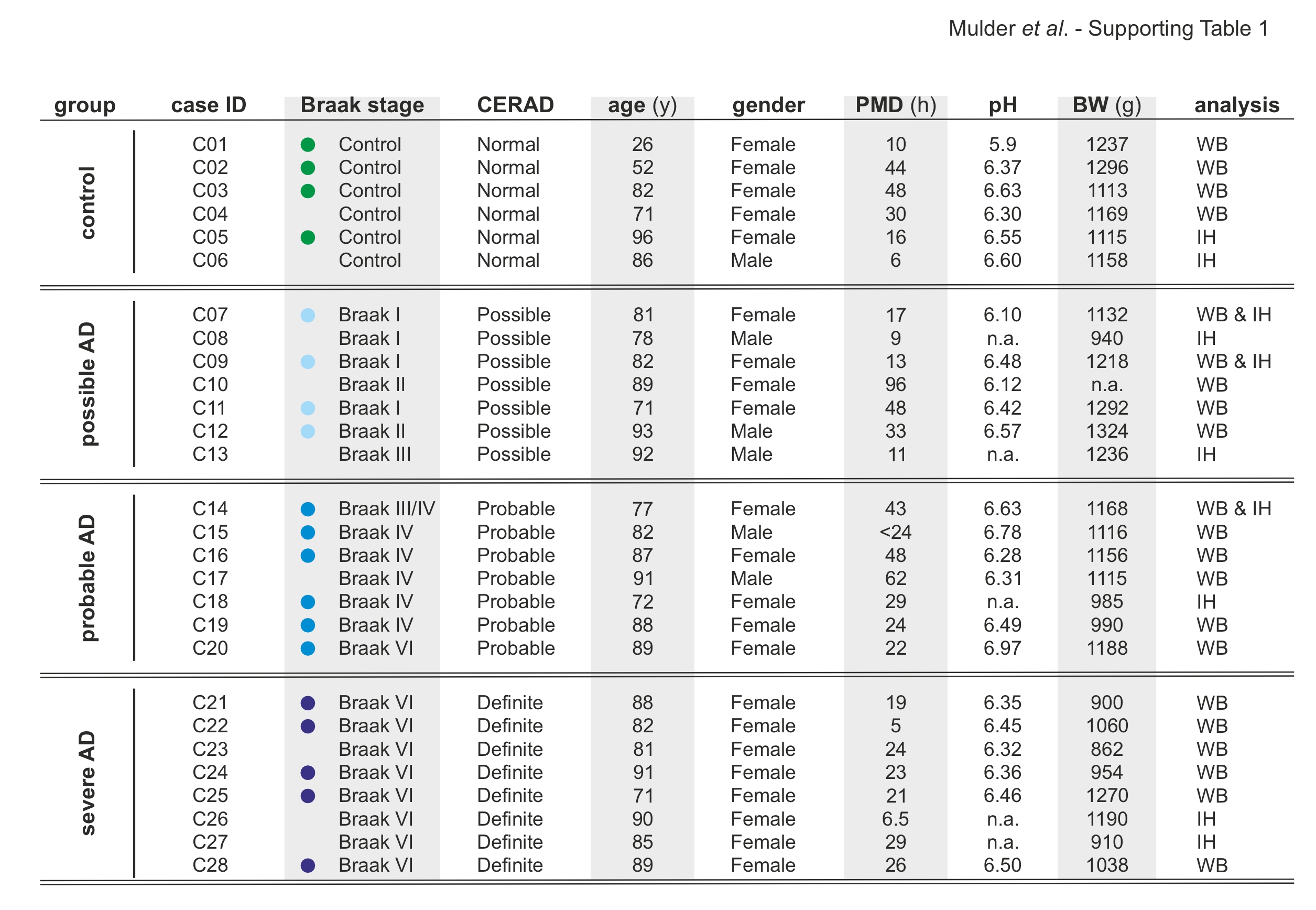

### Table S2

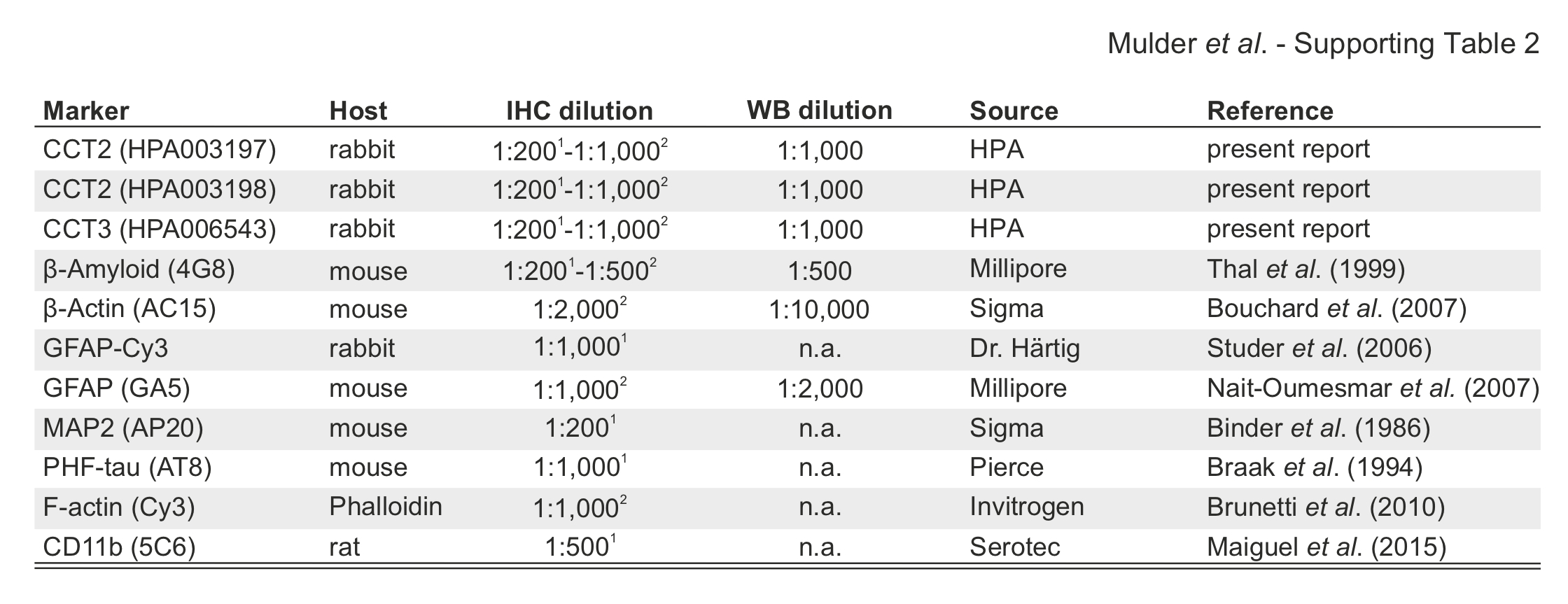
